# Bacteria facilitate viral co-infection of mammalian cells and promote genetic recombination

**DOI:** 10.1101/154021

**Authors:** A.K. Erickson, P.R. Jesudhasan, M.J. Mayer, A. Narbad, S.E. Winter, J.K. Pfeiffer

## Abstract

Intestinal bacteria promote infection of several mammalian enteric viruses, but the mechanisms and consequences are unclear. We screened a panel of 41 bacterial strains as a platform to determine how different bacteria impact enteric viruses. We found that most bacterial strains bound poliovirus, a model enteric virus. Given that each bacterium bound multiple virions, we hypothesized that bacteria may deliver multiple viral genomes to a mammalian cell even when very few virions are present, such as during the first replication cycle after inter-host transmission. We found that exposure to certain bacterial strains increased viral co-infection even when the ratio of virus to host cells was low. Bacteria-mediated viral co-infection correlated with bacterial adherence to cells. Importantly, bacterial strains that induced viral co-infection facilitated viral fitness restoration through genetic recombination. Thus, bacteria-virus interactions may increase viral fitness through viral recombination at initial sites of infection, potentially limiting abortive infections.

## INTRODUCTION

Enteric viruses, including poliovirus, reovirus, and norovirus, are spread through the fecal-oral route and initially replicate in the gastrointestinal tract where they encounter numerous resident bacteria (Pfeiffer and Virgin, 2016). Previously we and others demonstrated that gut microbiota promote replication, transmission, and pathogenesis of several enteric viruses (Baldridge et al., 2015; Jones et al., 2014; Kane et al., 2011; Kernbauer and Cadwell, 2014; Kernbauer et al., 2014; Kuss et al., 2011; Robinson et al., 2014; Uchiyama et al., 2014). Microbiota enhance replication and transmission of enteric viruses through several mechanisms (Karst, 2016; Pfeiffer and Virgin, 2016; Wilks et al., 2013). For example, microbiota can dampen host innate immune responses (Baldridge et al., 2015; Kane et al., 2011), or increase infectivity of viral particles by aiding attachment to host cells (Jones et al., 2014; Kuss et al., 2011; Robinson et al., 2014), or enhancing virion stability (Jones et al., 2014; Kuss et al., 2011; Li et al., 2015; Robinson et al., 2014). A poliovirus mutant with reduced bacterial binding had a transmission defect in mice due to virion instability in feces; therefore, virion-bacterial interactions likely promote transmission (Robinson et al., 2014).

Enteric viruses can bind to bacteria via bacterial surface polysaccharides. For example, human norovirus is thought to bind certain strains of bacteria by interacting with histo-blood group antigen glycans (Almand et al., 2017; Jones et al., 2014; Li et al., 2015; Miura et al., 2013). Poliovirus binds to bacterial *N*-acetylglucosamine-containing polysaccharides including lipopolysaccharide and peptidoglycan (Kuss et al., 2011; Robinson et al., 2014). Recently it was demonstrated that human norovirus can bind to different intestinal bacterial strains and multiple virions bound to a single bacterium (Almand et al., 2017; Li et al., 2015; Miura et al., 2013). It is unclear whether different bacterial strains bind to enteric viruses with different efficiencies. Furthermore, the consequences of virus-bacterial interactions are not completely understood.

RNA viruses such as poliovirus, reovirus, and norovirus exist as populations of genetically diverse viruses with varying levels of fitness (Domingo and Holland, 1997). Viral genetic diversity is generated through error-prone RNA replication. Mutations can have several consequences: most mutations are deleterious, some mutations are neutral, and a few mutations may be beneficial. Fitness of viruses with deleterious mutations can sometimes be restored by replication under high multiplicity of infection (MOI) conditions, which can facilitate genetic processes such as complementation and recombination (Domingo and Holland, 1997; Duarte et al., 1993; Duarte et al., 1994a; Muller, 1964; Nee, 1988). Mouse models of poliovirus infection have shown that both mutation and genetic recombination promote infection by driving viral adaptation necessary for viral replication and dissemination (Pfeiffer and Kirkegaard, 2005; Vignuzzi et al., 2006; Xiao et al., 2016). Poliovirus RNA recombination occurs in cells infected at high MOI (Egger and Bienz, 2002; Jarvis and Kirkegaard, 1992; Kirkegaard and Baltimore, 1986; Lowry et al., 2014; Runckel et al., 2013) and also occurs in the human gut after oral polio vaccination (Cuervo et al., 2001; Guillot et al., 2000; Minor et al., 1986). A basal requirement for genetic recombination is co-infection of a cell with at least two viruses. Co-infection of a cell with two or more viruses is unlikely when there are a limited numbers of viral particles, such as during the first cycle of replication following inter-host transmission. Recently it was demonstrated that poliovirus can spread as one unit containing multiple viral particles, either within lipid vesicles or as viral aggregates, and this delivery mode increased co-infection frequency and viral infectivity (Aguilera et al., 2017; Chen et al., 2015). How enteric RNA viruses generate high levels of population diversity upon the primary replication cycle within the host intestinal tract when a limited number of virions are present is unclear, but bacteria-mediated delivery of multiple virions is an intriguing possibility.

Here we use a diverse panel of bacterial strains as well as isogenic mutant strains to evaluate enteric virus-bacteria interactions and the consequences of these interactions on viral infection and evolution. By screening numerous bacterial strains we discovered that poliovirus-bacteria interactions vary among different bacteria. We found a large range of poliovirus-bacteria binding efficiencies, and we observed multiple poliovirions bound to the bacterial surface. Several bacterial strains enhanced viral co-infection efficiency and genetic recombination, even when the ratio of virions to host cells was low. Importantly, bacteria-mediated viral co-infection and genetic recombination generated variants that could replicate under conditions where the parental viruses could not. These results suggest that bacteria may increase viral fitness and population diversity.

## Results

### Screening Bacterial Strains for Poliovirus Binding

In order to investigate bacteria-virus interactions and effects on infection, we sought a diverse collection of bacterial species to screen a variety of parameters. In addition to 16 strains obtained from collaborators and 2 strains obtained from the American Type Culture Collection, we cultured 23 bacterial strains from the cecal contents of healthy mice using aerobic or anaerobic conditions. We obtained pure cultures and identified strains using 16S ribosomal DNA sequencing (Table S1)(Duerkop et al., 2012; Koropatkin et al., 2008; Stojiljkovic et al., 1995; Winter et al., 2010). In total, our collection includes 26 Gram-positive and 15 Gram-negative bacterial strains.

Our previous work indicated that poliovirus binds to bacteria via bacterial surface polysaccharides; therefore, we examined poliovirus-bacterial interactions using two assays. First, we evaluated the ability of bacteria to bind to poliovirus using electron microscopy (EM). EM analysis revealed that poliovirus bound to a variety of Gram-positive and Gram-negative bacteria. Additionally, we observed multiple virions bound to the surface of each bacterium (Fig. 1A-D). Second, we quantified the efficiency of poliovirus-bacteria binding using a pull-down assay. ^35^S-labeled poliovirus was incubated with bacteria for one hour prior to centrifugation, washing, and scintillation counting to determine the percent of input virus bound to bacteria. The level of non-specific viral binding was determined using inert beads with a diameter similar to bacteria. We found that 2% of input virus bound to beads, and 32 of the 36 bacterial strains initially tested bound significantly more virus as compared to beads (Fig. 1E). Poliovirus bound to both Gram-negative bacteria and Gram-positive bacteria, with binding efficiencies ranging from 4-37% of input virus bound (Fig. 1E). *Lactobacillus johnsonii,* a Gram-positive strain isolated from feces, bound poliovirus with the highest efficiency.

**Figure 1.**
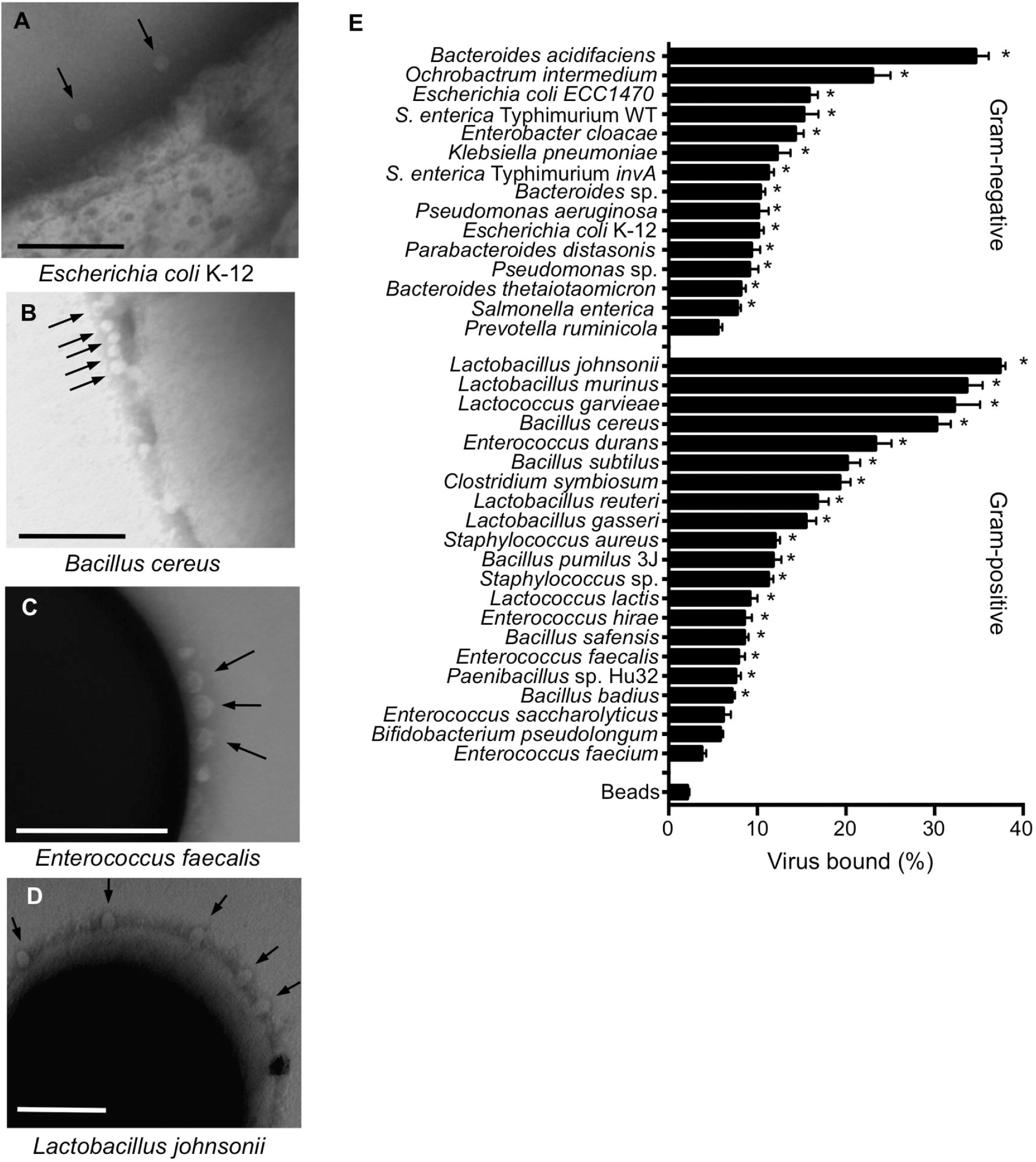
Poliovirus Binding to Bacterial Strains. Bacterial strains were either isolated from mouse cecal contents in this study (n=22) or were acquired from collaborators or the ATCC (n=14)(Table S1). (A-D) Electron micrographs of poliovirus bound to the surface of bacteria: 1x10^6^ CFUs of *Escherichia coli* K-12 (A), *Bacillus cereus* (B), *Enterococcus faecalis* (C), and *Lactobacillus johnsonii*- Fecal isolate (D) were incubated with 1x10^7^ PFU of purified poliovirus for 1 h prior to fixation with glutaraldehyde, staining with phosphotungstic acid, and imaging by TEM. Arrows are pointing to poliovirus virions on the bacterial surface. Scale bars represent 200 nm. (E) Pull down assay. 1x10^6^ PFU/5,000 CPM of ^35^S-labeled poliovirus was incubated with 1x10^9^ CFU of bacteria or inert beads for 1 h prior to centrifugation, washing, and scintillation counting of bacteria-associated ^35^S. Data are represented as mean ± SEM of the percent of input virus bound to the bacterial pellet. *p<0.05 versus Beads (one-way ANOVA followed by Dunnet’s multiple comparison test).

While working with *L. johnsonii*, we noticed that after 3-5 passages in liquid culture, the strain had altered properties suggestive of lab adaptation. For example, the original fecal isolate of *L. johnsonii* (called *L. johnsonii*-Fecal isolate) auto-aggregated following resuspension of bacterial pellets in PBS whereas the lab-adapted strain (called *L. johnsonii*-Lab passaged) did not (Fig. 2A). Lab adaptation of *L. johnsonii* strains can be conferred by changes in the synthesis of exopolysaccharide (EPS)(Horn et al., 2013). Previous studies demonstrated that *L. johnsonii* strains with mutations in the *eps* gene cluster have altered EPS production, auto-aggregation, and adhesion to host cells (Dertli et al., 2016; Dertli et al., 2015; Horn et al., 2013). Therefore, we examined EPS production in our *L. johnsonii*-Fecal isolate and *L. johnsonii*-Lab passaged strains compared to *eps* mutant strains using EM and a biochemical assay. Our collection of previously published isogenic *L. johnsonii* strains includes WT (FI9785, EPS positive, producing glycan EPS1 and heteropolysaccharide EPS2), *ΔepsA* (EPS negative), *epsC^D^*^88*N*^ (EPS overproducer), *ΔepsE* (reduced EPS with no EPS2 production)(Dertli et al., 2013; Dertli et al., 2016; Horn et al., 2013). First, EM analysis showed EPS on the cell surface of WT, Fecal isolate, and Lab passaged *L. johnsonii* strains (Fig. 2B, white arrow). Conversely EPS was absent from the *ΔepsA* mutant as previously reported (Fig. 2B)(Dertli et al., 2016). The EPS of the Lab passaged strain appeared slightly thicker and less defined as compared to the other strains (Fig. 2B, distance between white and black arrows). Second, we quantified purified EPS from the strains using a biochemical assay. Confirming previous findings (Dertli et al., 2016; Horn et al., 2013), *epsC^D^*^88*N*^ (EPS overproducer) had increased EPS production, accumulating up to 153% of WT. Conversely, EPS production by *ΔepsA* and *ΔepsE* was reduced compared to WT (Fig. 2C). We determined that the *L. johnsonii*-Fecal isolate and *L. johnsonii*-Lab passaged strains had similar EPS amounts to each other, which was lower but not significantly different from the amount determined for WT (Fig. 2C).

**Figure 2.**
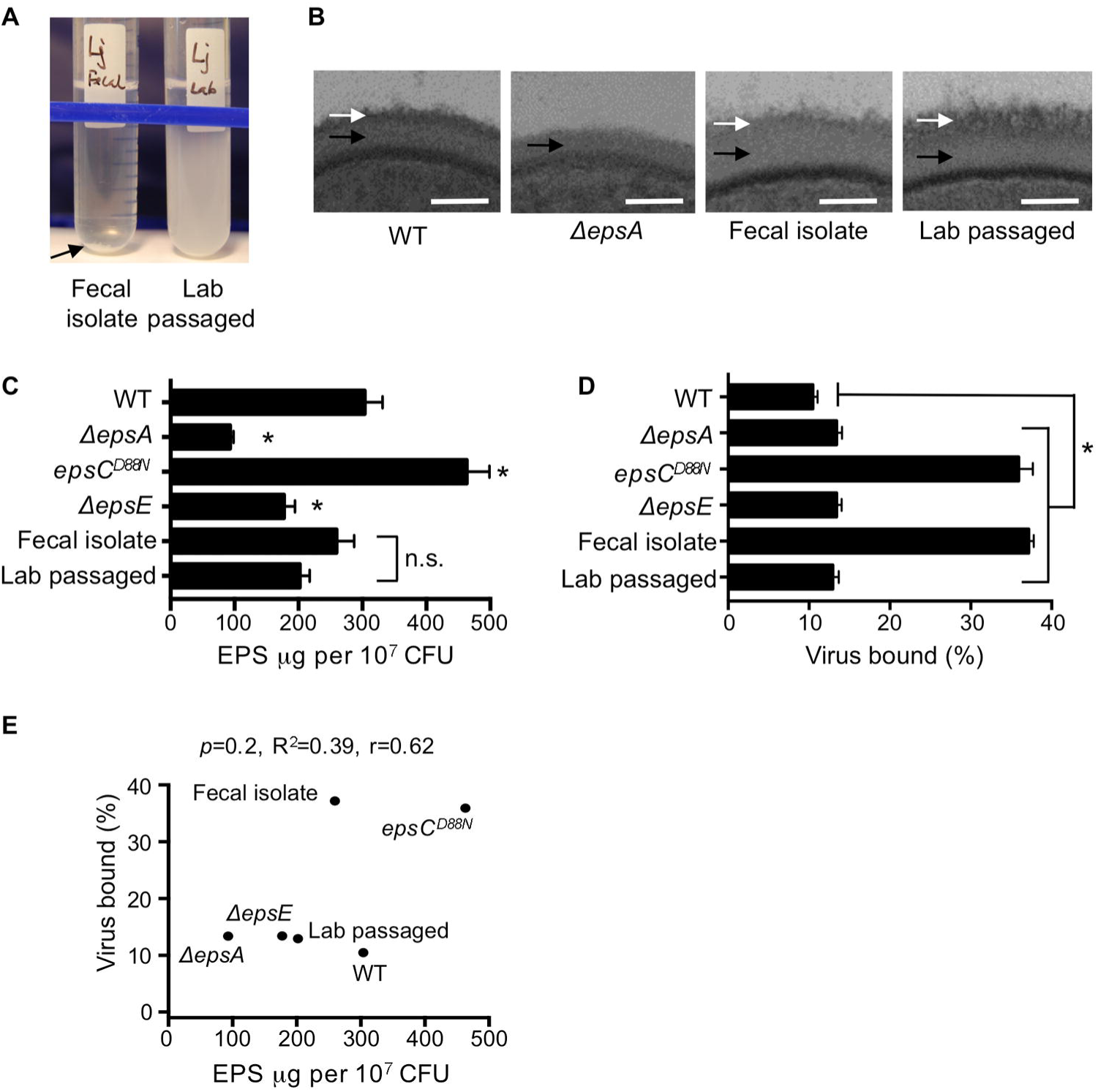
Exopolysaccharide (EPS) Production and Poliovirus Binding Efficiency of *Lactobacillus johnsonii* Strains. Several *L. johnsonii* strains were used in this work, including: Fecal isolate (isolated from mouse cecal contents), Lab passaged (a version of the Fecal isolate that was serially passaged in liquid media; a lab-adapted strain), WT (FI9785, a strain isolated from poultry)(Horn et al., 2013), *ΔepsA* (FI10938, WT strain lacking *epsA*, produces no EPS)(Dertli et al., 2016), *epsC^D^*^88*N*^ (FI10386, WT strain with a mutation in the putative chain length and polymerization protein EpsC, overproduces EPS and alters phenotype)(Horn et al., 2013), *ΔepsE* (FI10844, WT strain lacking *epsE*, produces altered/reduced EPS)(Horn et al., 2013). (A) Serial laboratory passage alters autoaggregation of the *L. johnsonii*- Fecal isolate. Overnight cultures of the *L. johnsonii*- Fecal isolate or *L. johnsonii*- Lab passaged strains were resuspended in PBS and were photographed after 5 min. Arrow indicates precipitated/autoaggregated cells for the Fecal isolate, but not the Lab passaged strain. (B) EM analysis of *L. johnsonii* strains comparing the EPS layer on the cell surface. Black arrows indicate the cell wall and white arrows indicate the EPS layer. Scale bars represent 25 nm. (C) Quantification of EPS from *L. johnsonii* strains. EPS was quantified using the phenol-sulfuric acid method with glucose as the standard. EPS quantities are shown as *μ* g per 10^7^ CFU. Data are represented as mean ± SEM (n=18, 3 independent experiments). \**p*<0.05 versus WT (ANOVA followed by Tukey’s post hoc test). (D) Quantification of poliovirus binding by *L. johnsonii* strains using the bacterial pull-down assay described in Figure 1E. Data are represented as mean ± SEM (n≥14, ≥3 independent experiments). \**p*<0.05 versus Beads (one-way ANOVA followed by Dunnet’s multiple comparison test). (E) Scatter plot of EPS (x-axis) versus percentage of virus bound (y-axis) for each *L. johnsonii* strain. Data points are the mean values presented in Figures 2C and 2D. *p* =0.2, R^2^=0.39, r=0.6206 (Pearson’s correlation coefficient calculation).

Since we previously found that bacterial surface polysaccharides bind poliovirus (Kuss et al., 2011; Robinson et al., 2014), we asked whether *L. johnsonii* EPS production impacts poliovirus binding using the pull-down assay described in Figure 1. We determined that the WT, *ΔepsA, ΔepsE*, and Lab passaged *L. johnsonii* strains had relatively low amounts (10-13%) of poliovirus bound, while the *epsC^D^*^88*N*^ and Fecal isolate strains had high amounts (>36%) of poliovirus bound (Fig. 1D). Notably, there was no correlation between EPS amounts and viral binding (Fig. 2E), suggesting that the ability of *L. johnsonii* to bind poliovirus is not dependent on the amount of EPS produced and that other factors contribute to the efficiency of viral binding.

### Certain Bacterial Strains Increase Viral Infectivity

Previously we found that exposure to bacteria or bacterial polysaccharides enhanced poliovirus infectivity in plaque assays (Kuss et al., 2011); therefore, we screened our panel of bacterial strains to determine whether strains differ in their capacity to promote poliovirus infection of cells. We used a high-throughput flow cytometry-based assay with polioviruses encoding DsRed or GFP (Teterina et al., 2010) to identify bacterial strains that enhance viral infection of HeLa cells (Fig. 3A). We used this assay previously to quantify viral co-infection frequencies from virion aggregates (Aguilera et al., 2017). To establish assay conditions, we performed mock infections, single infections with 1x10^4^ PFU DsRed or GFP poliovirus, or mixed infections of 1x10^6^ HeLa cells (MOI of 0.01). At 16 hpi, cells were fixed and the number of uninfected, single infected, or dual infected cells was quantified by flow cytometry. The total percentage of cells infected was determined as the percentage of DsRed+ and GFP+ gated cells (Fig. 3AB, and Fig. S1A). As predicted based on the MOI of 0.01, ~1% of cells exposed to both viruses were infected, with ~0.5% GFP+ and ~0.5% DsRed+ (Fig. 3C and S1A). To determine whether different bacterial strains enhance poliovirus infection, we pre-incubated 1x10^4^ PFU of DsRed and GFP viruses with inert beads or 1x10^8^ CFU bacteria, followed by infection of 1x10^6^ HeLa cells and analysis by the flow cytometry assay. We found that 32% (13/41) of the bacterial strains screened increased poliovirus infectivity, with *Prevotella ruminicola, Bacillus cereus,* and *Clostridia symbiosum* increasing infection two-fold more than beads (Fig. 3C). Initially, we hypothesized that poliovirus-bacteria binding efficiency would correlate with the HeLa cell infection efficiency. However, there was no correlation between binding efficiency of poliovirus to bacteria and the total number of infected cells (Fig. 3D). We also examined whether our collection of *L. johnsonii* strains enhance poliovirus infection, but only *L. johnsonii*-Fecal isolate increased infection marginally (1.5-fold, p<0.05)(Fig. 3E). Again, there was no significant correlation between the amount of virus bound to bacteria and the number of cells infected (Fig. 3F), suggesting that other factors such as the kinetics or strength for virus binding to bacteria may influence bacteria-mediated enhancement of poliovirus infection.

**Figure 3.**
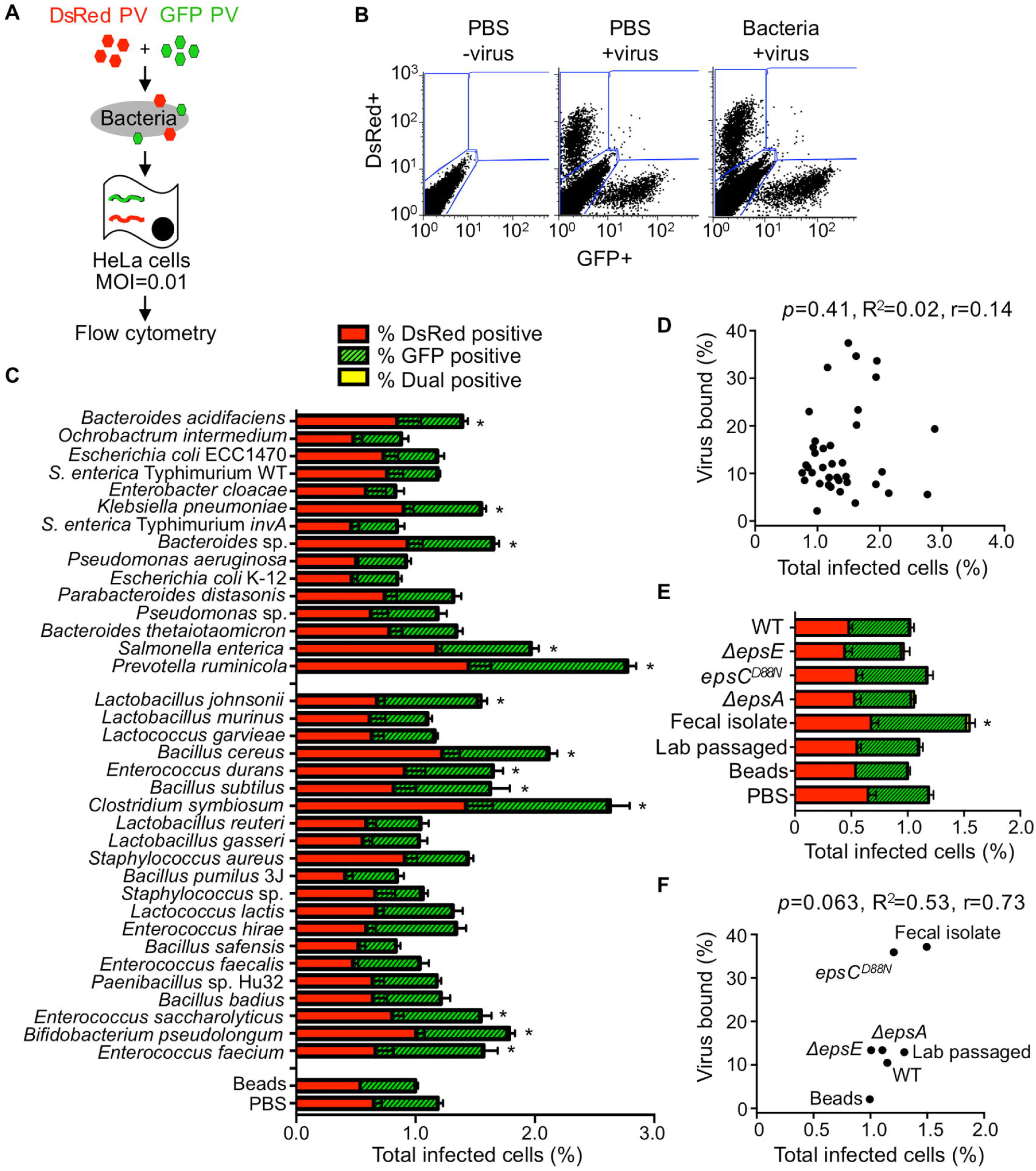
The Impact of Bacterial Strains on Viral Infectivity. (A) Schematic of flow cytometry-based infectivity assay that applies to Figures 3 and 4. 1x10^4^ DsRed- and/or GFP-expressing polioviruses were incubated with or without 1x10^8^ CFU bacteria for 1 h at 37°C prior to infection of 1x10^6^ HeLa cells (MOI of 0.01), bacteria and remaining virus were removed from cells by washing, infection proceeded for a single cycle (16 h), and DsRed and GFP positive cells were quantified by flow cytometry.(B) Representative FACs plots showing quantification of DsRed (top left gate), GFP (bottom right gate), and dual-infected cells (top right gate). Axes are fluorescence intensity of GFP (x-axis) and DsRed (y-axis). Each experiment counted ≥5x10^5^ events. (C) Total percentage of infected cells. Bars indicate the total percent of infected cells, which includes DsRed+ cells, GFP+ cells, or dual-positive cells. Bars are shaded to show the individual percentages of DsRed, GFP, or DsRed and GFP (dual) positive cells. Data are represented as mean± SEM (n=8-26). \**p*<0.05 versus PBS (one-way ANOVA followed by Dunnet’s multiple comparison test). (D) Scatter plot for correlation of total percent of infected cells and the percentage of virus bound for each bacterial strain. Data points are the mean values presented in Figures 3C and 1E. *p*=0.41, R^2^=0.02, r=0.14 (Pearson’s correlation coefficient). (E) The total percent of infected cells after viral incubation with or without *L. johnsonii* strains. Data are represented as mean ± SEM, n≥8. \**p*<0.05 versus PBS (one-way ANOVA followed by Dunnet’s multiple comparison test). (F) Scatter plot to test for correlation of total percent of infected cells and the percentage of virus bound for each *L. johnsonii* strain. Data points are the mean values presented in Figures 3E and 2D. *p*=0.06, R^2^=0.53, r=0.73 (Pearson’s correlation coefficient). See also Figure S1.

**Figure 4.**
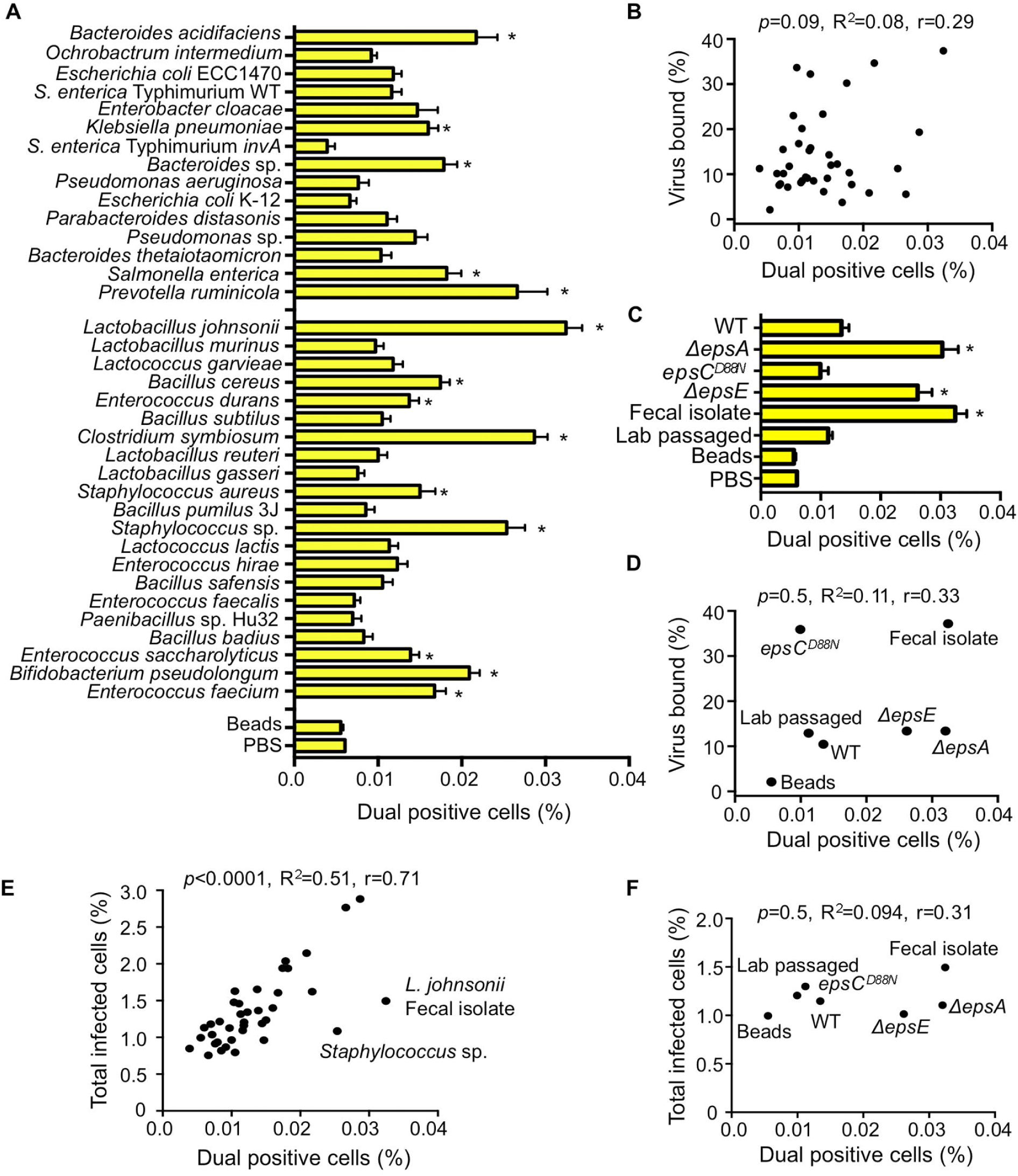
The Impact of Bacterial Strains on Viral Coinfection. (A) Percentage of dual infected cells positive for both DsRed and GFP determined using the flow cytometry-based assay described in Figure 3. Data are represented as mean ± SEM (n=8-52, ≥3 independent experiments). \**p*<0.05 versus Beads (one-way ANOVA followed by Dunnet’s multiple comparison test). (B) Scatter plot to test for correlation between the percentage of dual infected cells and the percentage of virus bound for each bacterial strain. Data points are the mean values presented in Figures 4A and 1E. *p*=0.1, R^2^=0.08, r=0.3 (Pearson’s correlation coefficient calculation). (C) Percentage of dual infected cells for *L. johnsonii* strains. Data are represented as mean ± SEM (n≥6). \**p*<0.05 versus Beads (one-way ANOVA followed by Dunnet’s multiple comparison test). (D) Scatter plot to test for correlation between the percentage of dual infected cells and the percentage of virus bound for each *L. johnsonii* strain. Data points are the mean values presented in Figures 4C and 2D. *p*=0.5, R^2^=0.1, r=0.3 (Pearson’s correlation coefficient calculation). (E) Scatter plot to test for correlation between the percentage of dual infected cells and the total percent of infected cells for each bacteria strain. Data points are the mean values presented in Figures 4A and 3C. *p*<0.0001, R^2^=0.5, r=0.71 (Pearson’s correlation coefficient calculation). (F) Scatter plot to test for correlation between the percentage of dual infected cells and the total percent of infected cells for each *L. johnsonii* strain. Data points are the mean values presented in Figures 4C and 3E. *p*=0.5, R^2^=0.1, r=0.31 (Pearson’s correlation coefficient calculation). See also Figure S1.

### Certain Bacterial Strains Facilitate Viral Co-infection

Since EM images in Figure 1 indicated that each bacterium binds multiple virions, we wondered whether bacteria facilitate delivery of more than one viral genome per HeLa cell, a process we refer to as bacteria-mediated viral co-infection. In particular, we hypothesized that bacteria may facilitate viral co-infection even when very little virus is present. To test this hypothesis, we infected 1x10^6^ HeLa cells with 1x10^4^ PFU of DsRed and GFP expressing polioviruses (MOI=0.01) with or without exposure to 1x10^8^ CFU of bacterial strains as described in Figure 3, and we quantified the number of cells infected with both viruses. We determined that GFP and DsRed viruses incubated with beads or PBS prior to infection of HeLa cells at an MOI of 0.01 resulted in approximately 0.006% of cells dual positive (GFP+DsRed+)(Fig. 4A and S1A), a value close to the predicted Poisson distribution value of 0.005%. We confirmed that dual-infected cells were from the initial viral replication cycle and not from secondary cycles of replication by demonstrating that the percentage of dual-infected cells at 16 hpi was not significantly changed in the presence of antibody blocking the poliovirus receptor to prevent secondary infection (Fig. S1B). We found that 39% of bacterial strains significantly enhanced the percent of dual-infected cells at least 2-fold over beads (Fig. 4A and 4C). Additionally, the *L. johnsonii*- Fecal isolate, *ΔepsA,* and *ΔepsE* strains all significantly enhanced dual infection by approximately 5-6 fold over beads (Fig. 4C). Similar to total infectivity, there was no correlation between viral binding efficiency and percentage of dual infected cells (Fig. 4B and 4D). However, there was a strong correlation between total infectivity and percent of dual infected cells (p<0.0001, Pearson r=0.72) (Fig. 4E). These results suggest that bacterial strains that increase infection efficiency also increase co-infection efficiency, likely because they effectively increase the MOI. However, four bacterial strains, *Staphylococcus* sp., *L. johnsonii*- Fecal isolate, *ΔepsA,* and *ΔepsE*, were outliers and did not display a linear relationship between infectivity and co-infection frequencies. Instead, for these four strains, the co-infection frequency was much higher than would be mathematically predicted from the total percent of infected cells (Fig. 4E-4F), suggesting additional factors play a role in bacterial mediated delivery of multiple viruses to a single cell.

### Bacteria-Mediated Viral Co-Infection Correlates with Bacterial Adherence to Host Cells

To further investigate the differential ability of bacterial strains to impact viral co-infection frequency we evaluated bacterial invasion of HeLa cells and adherence to HeLa cells. While viral binding efficiency to bacteria did not correlate with co-infection, we hypothesized that bacterial adherence to HeLa cells or invasion of HeLa cells may drive co-infection. To test this hypothesis, we began by measuring bacterial invasion using a gentamicin protection assay. For this assay, 1x10^6^ CFU of bacteria were incubated with 1x10^5^ HeLa cells for 1 h, treated with or without gentamicin for 2 h to kill extracellular bacteria, and the percent of intracellular bacteria was determined by CFU counts. Anaerobic bacterial strains were not included in these assays due to inconsistent survival through the aerobic experimental process. We found that wild-type *S. enterica* serovar Typhimurium, a positive control for invasion, had significantly higher numbers of intracellular bacteria compared to an isogenic invasion-deficient mutant (*invA* mutant)(Stojiljkovic et al., 1995; Winter et al., 2010)(Fig. 5A). All of the tested bacterial strains in our collection had less than 0.005% of input bacteria internalized, indicating that none are invasive (Fig. 5A and data not shown). To quantify bacterial adherence to HeLa cells, 1x10^6^ bacteria were incubated with 1x10^5^ HeLa cells for 1 h, followed by washing and quantification of cell-associated bacteria by CFU counts of HeLa lysate vs. input to reveal the percent of cell-associated bacteria. The adherence efficiencies of the bacterial strains varied greatly, ranging from 0.02 to 9% of cell-associated bacteria (Fig. 5B). To assess the relationship between adhesion efficiency of different bacteria strains and enhanced viral coinfection, the Pearson correlation coefficient was computed. A significant correlation was observed between the percent of cell-associated bacteria and the percentage of dual infected cells (p<0.0001, r=0.71, R^2^=0.5) (Fig. 5C). *L. johnsonii* strains were also screened for adhesion efficiencies. As expected from previous studies, we found that EPS levels of the isogenic *L. johnsonii* strains (Fig. 2C) inversely correlated with binding to mammalian cells (Fig. 5D)(Dertli et al., 2016; Horn et al., 2013). In contrast, despite having similar amounts of EPS (Fig. 2C), the *L. johnsonii*- Fecal isolate had 3-fold higher adherence to HeLa cells compared with the *L. johnsonii*- Lab passaged strain (Fig. 5D). For the *L. johnsonii* strains, the correlation between adhesion to HeLa cells and viral dual-infection was striking (p<0.0001, r=0.98, R^2^=0.96)(Fig. 5E). These results indicate that bacterial adhesion to HeLa cells may drive viral co-infection. Curiously, there was no correlation between the total percent of cells infected (DsRed, GFP, or both) and bacterial adhesion efficiency (Fig. S2A and S2B), suggesting that bacteria-mediated enhancement of infection efficiency is not sufficient to promote viral dual-infection. Collectively, these data suggest that enhanced ability of a bacterial strain to adhere to HeLa cells can greatly increase viral co-infection frequency in a manner independent of enhanced total infectivity.

**Figure 5.**
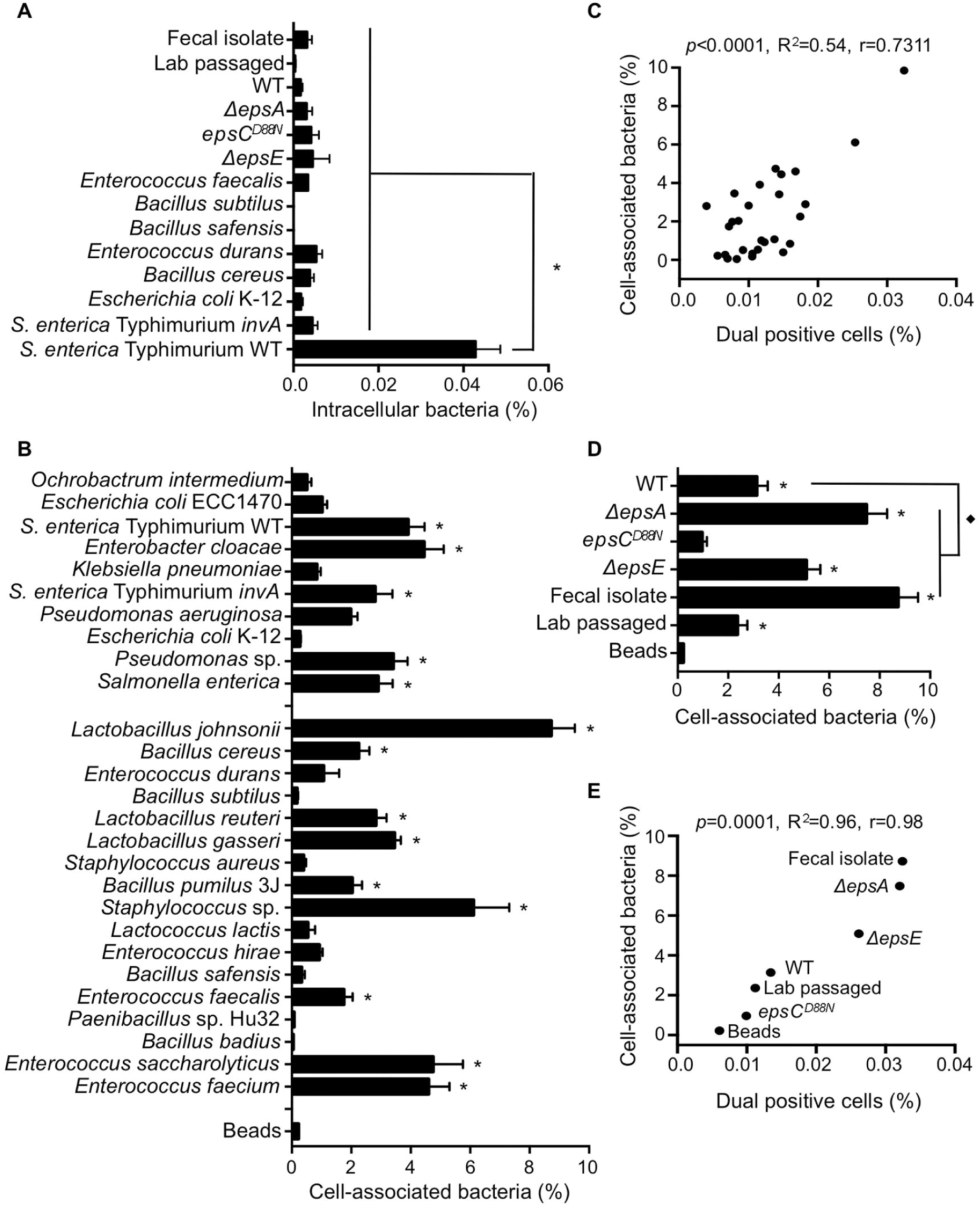
Bacterial Adherence to HeLa Cells and Impact on Viral Co-infection. (A) Bacterial invasion assay. 1x10^5^ HeLa cells were incubated with 1x10^6^ CFU bacteria for 1 h, washed, and treated with or without gentamicin to kill extracellular bacteria. HeLa cells were lysed and the number of intracellular bacteria determined from CFU counts, and data are reported as percentage of input CFU. Data are represented as mean ± SEM (n≥3). \**p*<0.0001 versus *S. enterica* serovar Typhimurium WT (one-way ANOVA followed by Dunnet’s multiple comparison test). (B) Bacterial attachment to HeLa cells. 1x10^5^ HeLa cells were incubated with 1x10^6^ CFU bacteria for 1 h, washed, and the number of cell-associated bacteria was enumerated by CFU counts in HeLa lysates. The percent of cell-associated bacteria is shown as the percentage of total input CFU. The percent of cell-association for the 2.8 *μ* m inert beads was determined from OD_600_ values of lysed Hela cells before (input) and after (attached) washing of the cell monolayers and are represented as percentage of input. Data are represented as mean ± SEM (n≥6). \**p*<0.05 versus beads (one-way ANOVA). (C) Scatter plot to test for correlation between the percentage of dual infected cells and the percentage of cell-associated bacteria for each bacterial strain. Data points are the mean values presented in Figures 4A and 5B. *p*<0.0001, R^2^=0.54, r=0.73 (Pearson’s correlation coefficient calculation). (D) Percentages of cell-associated bacteria for *L. johnsonii* strains. Data are represented as mean ± SEM (n≥6). \**p*<0.05 versus Beads or ♦*p*<0.05 versus WT (one-way ANOVA followed by Dunnet’s multiple comparison test). (E) Scatter plot to test for correlation between the percentage of dual infected cells and the percentage of cell-associated bacteria for *L. johnsonii* strains. Data points are the mean values presented in Figures 4C and 5D. *p*=0.0001, R^2^=0.96, r=0.98 (Pearson’s correlation coefficient calculation). See also Figure S2.

### Exposure to Certain Bacterial Strains Enhances Poliovirus Genetic Recombination

Several studies have suggested that RNA viruses benefit from the delivery of multiple viral genomes to a single cell (Aguilera et al., 2017; Chen et al., 2015; Combe et al., 2015; Domingo and Holland, 1997; Duarte et al., 1993; Duarte et al., 1994b; Novella et al., 1995). Therefore we asked whether bacterial strains with differential abilities to facilitate viral co-infection under low MOI conditions could affect the poliovirus genetic recombination frequency and restoration of viral fitness. To quantify poliovirus recombination frequencies, we employed a well-characterized system consisting of two viruses that can be discriminated from each other and from recombinants based on a simple phenotypic assay (Kirkegaard and Baltimore, 1986). Wild-type poliovirus is sensitive to the drug guanidine hydrochloride (Drug^S^), but is able to grow at the high temperature of 39.5°C (Temp^R^). Conversely, a virus with mutations in the 2C coding region and an insertion in the 3’ non-coding region is resistant to guanidine hydrochloride (Drug^R^) and temperature sensitive (Temp^S^). Upon high MOI infection with the Drug^S^/Temp^R^ and Drug^R^/Temp^S^ viruses, recombination can generate a variety of variants, including those that are Drug^R^/Temp^R^ and are able to replicate in the presence of drug at 39.5°C (Fig. 6A).

**Figure 6.**
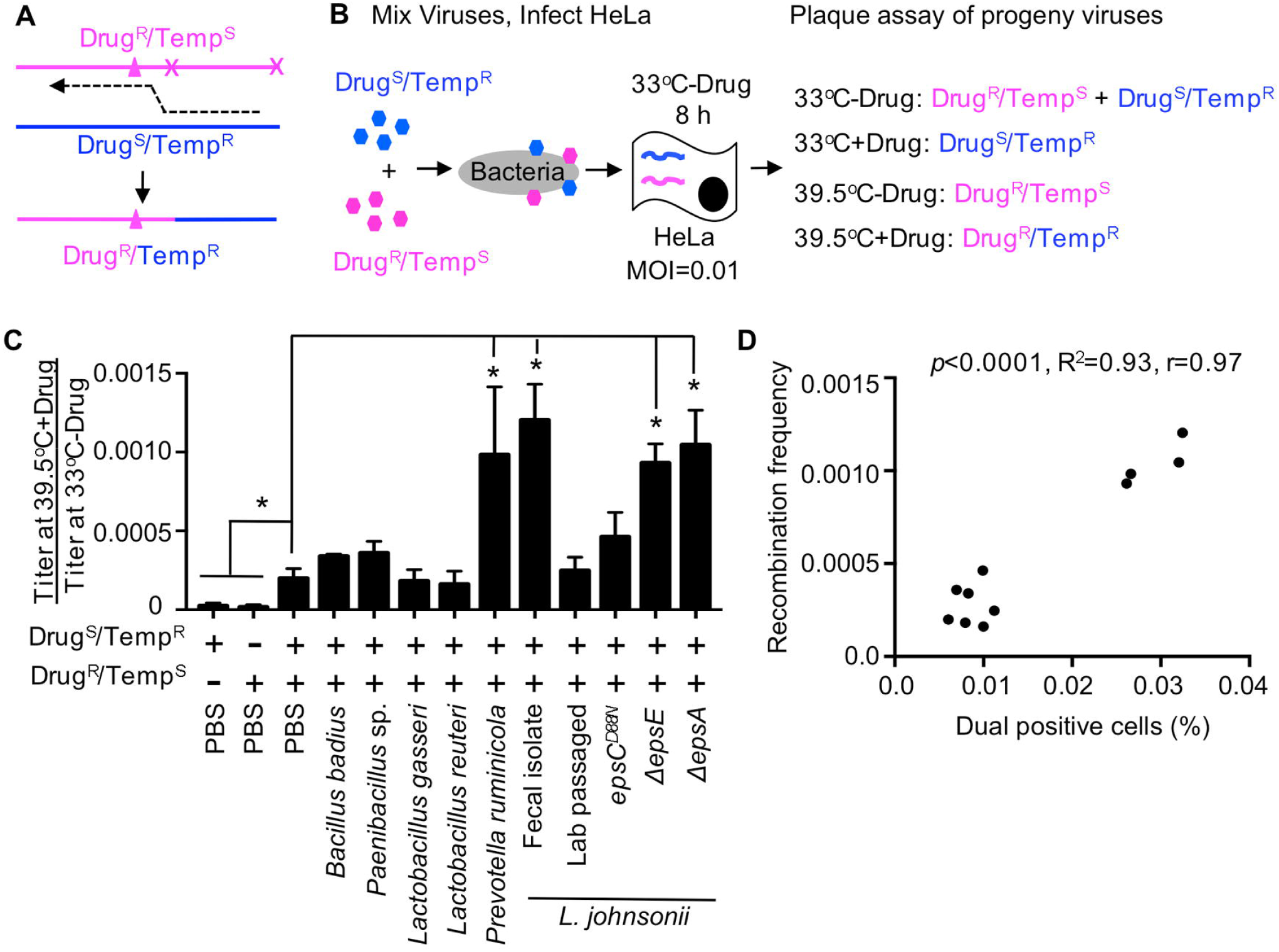
Effect of Bacterial Strains on Poliovirus Recombination Frequency. (A) Diagram of recombination between Drug^R^/Temp^S^ and Drug^S^/Temp^R^ parental polioviruses. Triangle denotes mutation conferring guanidine (Drug) resistance and **x** denotes site of temperature sensitive mutations inhibiting replication at 39.5°C (Kirkegaard and Baltimore, 1986). The dotted line indicates the location of a recombination event that creates progeny that are able to grow at 39.5°C in the presence of 1 mM guanidine (Drug^R^/Temp^R^). (B) Schematic of recombination assay. 1x10^5^ PFU of each parental virus was mixed and incubated with 1x10^8^ CFU bacteria prior to infection of 1x10^7^ HeLa cells (MOI of 0.01). Infection was allowed to proceed for 8 h under permissive conditions for both viruses (33°C without drug) and progeny viruses were quantified by plaque assay at both permissive and restrictive conditions to determine yields of individual parental viruses and recombinants. (C) Recombination frequencies after exposure to bacterial strains. Recombination frequencies are shown as the viral titers (PFU/mL) at 39.5°C+Drug divided by viral titers at 33°C-Drug. HeLa cells were also infected in parallel with each parental virus alone as controls to calculate the frequency of reversion and de novo mutation acquisition. Data are represented as the mean ± SEM (n=4-12), from at least 2 independent experiments. \**p*<0.01 versus the mix of parental viruses in PBS (Student’s unpaired t test). (D) Scatter plot to test for correlation between the percentage of dual infected cells and the recombination frequency for each bacterial strain. Data points are the mean values presented in Figures 6C, 4A and 4C. *p*<0.0001, R^2^=0.93, r=0.97 (Pearson’s correlation coefficient calculation). See also Figure S3.

To determine whether bacteria can influence genetic recombination even under low MOI conditions, we incubated 1x10^5^ PFU of both the Drug^S^/Temp^R^ and Drug^R^/Temp^S^ parental viruses or each alone with PBS or 1x10^8^ CFU bacteria prior to infection of 1x10^7^ HeLa cells (MOI=0.01) under dual permissive conditions (33°C, no drug) to facilitate replication of both parental viruses. At 8 hpi, viral progeny were harvested and quantified by plaque assays performed under permissive and restrictive growth conditions to calculate recombination frequencies and frequencies of confounding variables such as reversion of Drug^R^ or Temp^S^ mutations or *de novo* mutation acquisition (Fig. 6B and S3A-S3B). We found that each parental virus produced progeny with the expected frequencies/phenotypes during single infections: Both replicated efficiently at 33°C-drug, the Drug^S^/Temp^R^ parent had a 6,200-fold replication defect at 33°C+drug, the Drug^R^/Temp^S^ parent had a 26,000-fold replication defect at 39.5°C-drug, and both viruses had a >75,000-fold replication defect at 39.5°C+drug (Fig. S3B). However, when both viruses were incubated together in PBS prior to infection of HeLa cells, a small number of plaques formed at 39.5°C+drug (Fig. S3AB). This frequency of plaques at 39.5°C+drug was 20-fold higher than that observed from either parental virus alone, indicating that reversion or *de novo* Drug^R^ or Temp^R^ mutation acquisition was negligible for these assays. By dividing the viral yield at 39.5°C+drug by the viral yield at 33°C-drug, we determined that viruses incubated with PBS had an observed recombination frequency of 2×10^−4^. When we incubated the parental viruses with bacteria prior to the infection, we found that recombination frequencies ranged from 1.6×10^−4^ to 1.2×10^−3^ (Fig. 6C). *Prevotella ruminicola*, *L. johnsonii*- Fecal isolate, *ΔepsE,* and *ΔepsA* strains all significantly increased recombinant yields by more than 4.6-fold over PBS (Fig. 6C and S3AC). We determined through sequence analysis that 83.3% of scored plaques were indeed genotypic recombinants, confirming the results from the phenotypic plaque assays (Fig. S3AD). Next, we assessed the relationship between recombination frequencies (Fig. 6C) and co-infection frequencies (Fig. 4A) for each bacterial strain and we observed a striking correlation (p<0.0001, r=0.97, R^2^=0.95)(Fig. 6D). These data together suggest that poliovirus interactions with certain bacterial strains prior to initial infection of host cells increases the probability that a cell will be infected with multiple viruses even at low viral concentrations. Furthermore, bacterial enhancement of viral co-infection can facilitate recombination between distinct viruses giving rise to progeny with the ability to grow under otherwise restrictive conditions.

## DISCUSSION

As a platform to understand how bacteria can impact enteric viruses, we screened several bacterial strains for the capacity to bind to poliovirus, facilitate viral infection and co-infection of mammalian cells, and promote viral genetic recombination. To mimic the first cycle of viral replication upon inter-host transmission, we used low MOI conditions in the presence or absence of bacteria. We found that exposure to certain bacterial strains enhanced viral infectivity and increased viral co-infection frequency even when the ratio of virus to host cells was low. Contrary to our initial hypothesis, viral binding efficiency to bacterial strains was not the primary determinant of bacteria-mediated viral co-infection. Instead, viral co-infection correlated with the ability of bacteria to adhere to mammalian cells. Importantly, bacteria-mediated viral co-infection had functional consequences: bacterial strains that induced viral co-infection facilitated fitness restoration through genetic recombination, with progeny viruses able to replicate under conditions where parental viruses could not. Thus, bacteria-virus interactions may facilitate viral propagation and adaptation at initial sites of infection within a host.

First, we found that poliovirus bound to different bacterial strains with different efficiencies. Notably, although 90% of bacterial strains bound to poliovirus more efficiently than inert beads, only 5 of the 41 bacterial strains bound more than 30% of viral input, including one Gram-negative strain (*Bacteroides acidifaciens*) and four Gram-positive strains (*Bacillus cereus*, *Lactococcus garvieae*, *Lactobacillus murinus*, and *Lactobacillus johnsonii*). A specific poliovirus-bacteria binding ligand or common factor shared among these high binding strains has not been identified. However, poliovirus binding with high efficiency to few bacterial strains suggests some specificity of the bacteria-poliovirus interaction. Miura *et al.* demonstrated that *Enterobacter cloacae* binds to human norovirus particles through histo-blood group antigen-like molecules localized in the extracellular polymeric substances of the bacteria (Miura et al., 2013). To determine if bacterial exopolysaccharide production plays a role in poliovirus-bacterial interactions we compared the high poliovirus-binding *L. johnsonii*- Fecal isolate with a panel of isogenic *L. johnsonii* strains that have altered EPS production and structure (Dertli et al., 2013; Dertli et al., 2016; Horn et al., 2013). We found that total EPS amounts produced by the *L. johnsonii* strains did not correlate with poliovirus binding or infectivity (Fig. 2 and 3E). It is possible that binding is associated with specific qualities of the EPS such as composition or chain length rather than quantity (Horn et al 2013), or with other surface structures. While specific bacterial ligands important for poliovirus binding to these strains remain unclear, we were able to use this collection of bacterial strains and the *L. johnsonii* isogenic mutants to examine factors that impact bacteria-mediated viral infection and co-infection.

Next, we used a flow-cytometry based assay to determine whether the bacterial strains enhance poliovirus infectivity and co-infection of cells when the ratio of viruses to mammalian cells is low. Using a panel of four bacterial strains, our previous work demonstrated that bacteria were capable of increasing poliovirus infectivity in plaque assays (Kuss et al., 2011), but whether the infectivity enhancement extends to other strains was unknown. Here, we determined that 13 of the 41 bacterial strains increased viral infectivity using the flow-cytometry based assay (Fig. 3C and 3E). Because only 32% of bacterial strains increased viral infectivity in this assay, bacteria differ in their capacities to promote viral infection.

EM images showed multiple virions bound to various bacterial strains (Fig. 1A-1D); therefore, we hypothesized that bacteria may deliver more than one viral genome to a mammalian cell. To test this hypothesis, we used the flow-cytometry based assay to quantify the percentage of cells infected with both a DsRed- and a GFP-expressing virus. The percentage of dual infected cells for polioviruses incubated with inert beads or PBS was close to the predicted Poisson distribution value (1% of cells infected; 0.005% of cells dual infected at MOI of 0.01). However, the actual co-infection frequency is likely higher than that observed here since cells infected with two DsRed or two GFP viruses cannot be scored in this assay. We found that 39% of the bacterial strains increased poliovirus co-infection frequency (Fig. 4A and 4C). Not surprisingly, there was a strong correlation between co-infection frequency and the total percent of infected cells for most bacteria strains (Fig. 4E). However, four strains (*Staphylococcus* sp., *L. johnsonii*- Fecal isolate, *ΔepsE*, and *ΔepsA*) were outliers and increased co-infection far more than would be predicted from the total number of infected cells (Fig. 4E-4F). We do not know why these four Gram-positive strains increased viral co-infection without increasing total infection to a similar extent. It is possible that these strains deliver a relatively large number of virions to a relatively small number of cells, and that infection events are underestimated in the assay since cells infected with two or more DsRed or GRP viruses are not discriminated from singly infected cells. Indeed, EM images of *L. johnsonii*- Fecal isolate show dozens of viral particles on the bacterial surface (Fig. 1D and data not shown). While bacteria-mediated viral co-infection did not correlate with the viral binding efficiency to bacteria (Fig. 4B and 4D), it did correlate with bacterial adhesion to mammalian cells (Fig. 5C and 5E). These data suggest a model where bacteria must bind at least two virions (likely more), and adhere to cells to promote viral co-infection.

Because certain bacterial strains facilitated viral co-infection of cells, we wondered whether bacteria increase viral fitness by promoting genetic recombination. Viral recombination requires that two or more viruses co-infect the same cell. Therefore, previous experiments examining poliovirus recombination frequencies in cell culture were performed at high MOI (>10). Under high MOI conditions, several groups have observed recombination frequencies indicating that 1-20% of single cycle progeny viruses are recombinants (Jarvis and Kirkegaard, 1992; King, 1988; Kirkegaard and Baltimore, 1986; Lowry et al., 2014; Runckel et al., 2013). Because the first cycle of replication following inter-host transmission is likely initiated by a small number of virions at mucosal sites, we sought to determine whether recombination is detectable under low MOI conditions and whether bacteria can facilitate viral genetic recombination by delivering more than one viral genome per cell. We hypothesized that, at an MOI of 0.01, recombinants should be rare unless bacteria facilitate synchronous viral co-infection. Polioviruses with distinct genetic defects that restrict viral growth under certain conditions were incubated with selected bacteria strains that displayed a range of co-infection frequencies (Fig. 4A). The frequency of recombinant progeny was determined after one round of replication performed under permissive conditions. We determined that PBS-treated parental viruses generated a recombination frequency of 2x10^−4^ (for the 190 nt interval between relevant alleles) under MOI 0.01 conditions. This recombination frequency was much higher than expected, given that previous work with these viruses using an MOI of 60 generated a recombination frequency of 1.3x10^−3^ (Kirkegaard and Baltimore, 1986). It is possible that recombination frequency does not correlate linearly with MOI, that the later time point for progeny collection used here (8 hpi vs. 4 hpi) enhances the observed recombination frequency, or that viral aggregates facilitated co-infection as we have recently shown (Aguilera et al., 2017). While our observed recombination frequency for PBS-treated parental viruses was 2x10^−4^, bacteria-treated parental viruses generated recombination frequencies of 1.6x10^−4^ to 1.2x10^−3^. Recombinant progeny yields were significantly increased, up to 3.5-fold, for 4 out of 10 bacteria strains evaluated: *Prevotella ruminicola*, *L. johnsonii*- Fecal isolate, *ΔepsE*, and *ΔepsA* strains (Fig. 6C). As predicted, we found a striking correlation between bacteria mediated co-infection and poliovirus recombination frequency (Fig. 6D).

Our results indicate that bacteria-mediated viral co-infection can facilitate viral recombination events with the potential to drive viral adaptation and increase viral fitness. While our model system uses defined genetic markers for ease of detection, our results would theoretically apply to escape from a variety of detrimental mutations present in a viral population. We speculate that bacteria-mediated enhancement of viral recombination in the first cycle of replication in a newly infected host may rescue viral defects that may otherwise culminate in abortive infection. Overall, by screening a large panel of bacterial strains, our work revealed new insights into virus-microbiota interactions and expands our understanding of how microbiota promote enteric viral infections.

## Methods

### Mice

Mice were handled according to the Guide for the Care of Laboratory Animals of the National Institutes of Health. All mouse studies were performed at UT Southwestern (Animal Welfare Assurance #A3472-01) using protocols approved by the local Institutional Animal Care and Use Committee. C57BL/6 *PVR-IFNAR*^−/−^ mice, expressing the human PVR and deficient for the IFN *α / β* receptor, were obtained from S. Koike (Tokyo, Japan)(Ida-Hosonuma et al., 2005). Six-week old male mice were euthanized and cecal contents were collected to isolate bacterial strains as described below.

### Bacterial Strains

Bacterial strains were either acquired from collaborators, the ATCC, or were isolated from the cecal contents of healthy mice, as indicated in Table S1. For fecal isolates, fresh feces were resuspended in PBS containing 0.1% L-Cysteine (as a reducing agent to aid survival of anaerobic bacteria) prior to dilution and plating on rich media (see Table S1). Plates were incubated at 37^o^C for 24-72 h in normal atmospheric conditions (aerobic), in incubators supplemented with 5% CO_2_, or in an anaerobic chamber. Well separated colonies were picked, streaked for isolation, and identified based on 16S rDNA sequencing of PCR products generated with one of two universal primer sets (Forward: 5’ AGAGTTTGATYMTGGCTCAG, Reverse: 5’ ACGGYTACCTTGTTACGACTT or Forward: 5’ CCAGACTCCTACGGGAGGCAGC, Reverse: 5’CTTGTGCGGGCCCCCGTCAATTC (Rudi et al., 1997). For experiments, 25 mL overnight liquid cultures of bacterial strains inoculated from frozen glycerol stocks were incubated under conditions/media listed in Table S1. The next day, cultures were centrifuged at 4,000 rpm for 20 min, pellets were washed in PBS, and bacteria were resuspended in PBS. Bacterial concentrations were determined by OD_600_ values and comparison to standard curves of CFU dilutions.

### Viruses and Cells

Poliovirus work was performed under BSL2+ conditions as recommended by the World Health Organization. All experiments used serotype 1 Mahoney poliovirus and HeLa cells. HeLa cells were a gift from K. Kirkegaard and were authenticated in March, 2016 by the University of Arizona Genetics Core using short tandem repeat (STR) profiling with the Promega PowerPlex16HS Assay. HeLa cells were grown in Dulbecco’s modified Eagle’s medium (DMEM) supplemented with 10% calf serum and 1% penicillin-streptomycin in 5% CO_2_. Virus stocks were generated from infectious clone plasmids as previously described (Kuss et al., 2011). The Drug^R^/Temp^S^ virus, 3NC-202guaR, contains a mutation that confers guanidine resistance (2C-M187L) and two mutations that confer temperature sensitivity (2C-V250A and an insertion in the 3’ noncoding region, 3NC-202)(Kirkegaard and Baltimore, 1986; Sarnow et al., 1986). Wildtype poliovirus was used as the Drug^S^/Temp^R^ virus. DsRed- and GFP-expressing viruses contain fluorescent protein sequences inserted after amino acid 144 of poliovirus protein 2A (Teterina et al., 2010).

### Electron Microscopy

Poliovirus was purified by cesium chloride gradient centrifugation, followed by concentration and desalting using Amicon filters (Millipore) as previously described (Kuss et al., 2011). Prior to imaging, 1x10^6^ CFU (as determined by OD_600_ values) of indicated bacterial strains was incubated with or without 1x10^7^ PFU poliovirus for 1 h at 37^o^C to facilitate binding. Samples were fixed with 2.5% glutaraldehyde for 1 h at room temperature and 2.5 μl of the inactivated virus was placed on 400 mesh carbon-coated copper grids that had been glow discharged for 30 s using PELCO EasiGlowTM 91000. The grids were stained with 2% phosphotungstic acid and examined using a TEI Technai G^2^ Spirit Biotwin transmission electron microscope (FEI, Hillsboro, OR) equipped with Gatan ultrascan charge-coupled-device (CCD) camera, and with a side-mounted SIS Morada 11-megapixel CCD camera, operated at an acceleration voltage of 120 kV. Images were taken at magnifications of 43,000x and 60,000x.

### Viral Pull Down Assays

^35^S-labeled poliovirus (1x10^6^ PFU/5,000 CPM) was generated as previously described (Kuss et al., 2011) and incubated with 1x10^9^ CFU bacteria (as determined by OD_600_ values) or ~3.5x10^6^ inert beads (Dynabeads M-280 Streptavidin, Invitrogen) for 1 h at 37°C to facilitate viral binding. Following incubation, samples were centrifuged at 4000 rpm for 20 min, washed with PBS, and ^35^S was quantified in a scintillation counter.

### EPS Isolation and Quantification

Capsular EPS was isolated using methods adapted from (Dertli et al., 2016). *Lactobacillus* strains were inoculated from glycerol stocks into 20 mL of MRS broth and grown overnight, and these cultures were used to inoculate 500 mL of MRS broth, and cultures were grown for 48 h. Cultures were agitated vigorously to disperse aggregates and aliquots were plated for total CFU determination. Bacteria cells were pelleted by centrifugation at 6000 x g for 30 min at 4°C and washed twice. Bacterial pellets were resuspended in 50 mL of 1M NaCl and capsular EPS was extracted from cells by sonication (4 times for 45 seconds at 30% amplitude). To remove insoluble material samples were centrifuged at 6000 x g for 30 min and supernatants were collected. EPS was precipitated by adding an equal volume of ethanol and incubating at −20°C overnight, then centrifuged at 10,000 x g for 30 min at 4°C and supernatant was discarded. Pellets were resuspended in water and EPS was recovered with ethanol precipitation repeated twice, with final resuspension in 20 mL water, followed by dialysis for 72 h at 4°C using 12-14 kDa dialysis membranes. Samples from dialysis tubing were then lyophilized by freeze drying for 2 days. Freeze dried pellets were weighed and resuspended in 10 mL water to provide crude EPS for further analysis.

EPS produced from each of the *L. johnsonii* strains was quantified using the phenol-sulfuric method (DuBois et al., 1956; Masuko et al., 2005) with glucose as the standard. A standard curve was generated using D-Glucose (Sigma) in water. EPS samples were serially diluted in 100 μL water in glass tubes and 100 μL 5% phenol/water (Sigma-P9346) was added. Samples were vortexed and 500 μL of concentrated sulfuric acid (96%) was added, mixed, and 200 μL was used for absorbance readings at 490nm. EPS quantity of each sample was calculated using glucose standard curve and is expressed as μg per total CFU.

### Flow Cytometry

The flow cytometry based assay for co-infection was performed as previously described (Aguilera et al., 2017). Polioviruses were derived from Mahoney serotype 1 poliovirus with DsRed or GFP fluorescent proteins inserted after amino acid 144 of poliovirus protein 2A (Teterina et al., 2010). 1x10^4^ PFU of DsRed-Poliovirus and 1x10^4^ PFU GFP-Poliovirus were incubated in 200 μL PBS+ (PBS supplemented with 100 μg/mL MgCl_2_ and CaCl_2_) or 1x10^8^ CFU bacteria or Beads (Dynabeads M-280 Streptavidin, Invitrogen) for 1 h at 37°C. Samples (200 μL total) were added to 2x10^6^ HeLa cells for 30 min at 37°C. Cell monolayers were washed 3 times and 2 mL DMEM supplemented with 5% calf serum and 1% penicillin-streptomycin was added to cells. After incubation at 37°C for 16 h cells were harvested using 0.1% trypsin- 0.05% EDTA, washed and fixed with 2% paraformaldehyde/PBS for 15 min at room temperature, washed and resuspended in PBS containing 2% fetal bovine serum. DsRed and GFP expression was determined from at least 5x10^5^ events using a FACSCalibur cytometer equipped with 488- and 635-nm lasers. Data were analyzed using FlowJo software.

### Bacterial Invasion and Adherence Assays

Bacteria were inoculated from glycerol stocks into 5 mL growth medium and grown overnight as described above in Bacterial Strains and Table S1. 1mL of culture was centrifuged at 5,000 x g for 5 min, washed and resuspended in 1 mL PBS. The OD_600_ was measured and bacteria diluted in PBS to reach OD_600_=0.01, or approximately 1x10^6^ CFU bacteria in 100 μL PBS. Serial dilutions were plated on appropriate agar plates and incubated at 37°C for 24-72 h for input CFU/mL determination, or added to 10^5^ HeLa cells for experiments. After incubation at 37°C for 1 h cells were washed 5 times with PBS and 0.5 mL of DMEM and 5% calf serum (for attachment assay) or DMEM and 5% calf serum and 100 μg/mL Gentamicin (for invasion assay) was added. After incubation at 37°C for 2 h cell monolayers were washed 3 times with PBS and 0.5 mL 1% TritonX-100 in PBS was added for 15 min at room temperature to lyse HeLa cells. The lysate was collected and mixed vigorously. Serial dilutions of lysate in PBS was plated on appropriate agar plates and incubated at 37°C for 24-72 h. Bacterial colonies were counted and CFU/mL determined. We quantified the adherence of control beads (Dynabeads M-280 Streptavidin, Invitrogen) using OD_600_ analysis. The percent of cell-associated beads was calculated by the OD_600_ of beads+wash divided by OD_600_ of bead-wash.

### Recombination Assays

Viruses used for the recombination assay, wild type Mahoney serotype 1 poliovirus (Drug^S^/Temp^R^) and mutant poliovirus 3NC-202gua^R^ (Drug^R^/Temp^S^) were previously described (Kirkegaard and Baltimore, 1986). Prior to infection 1x10^5^ PFU of both the Drug^S^/Temp and Drug^R^/Temp^R^ parental viruses or 1x10^5^ PFU of each alone were incubated with 1 mL PBS or 1 mL PBS and 1x10^8^ CFU bacteria at 37°C for 1 h, then added to 1×10^7^ HeLa cells (MOI=0.01). After incubation at 33°C for 30 min, virus and bacteria were removed by washing 2 times with PBS. 10 mL of DMEM containing 5% calf serum and 1% penicillin-streptomycin was added and plates incubated at 33°C for 8 h. Cells were harvested by scraping and resuspended in 1 mL of PBS followed by freeze thawing three times to lyse the cells and release progeny viruses. Samples were centrifuged to remove cell debris at 1500 x g for 10 min and viral supernatant collected. Dilutions of the viral stocks were added to 4 x 6-well plates of HeLa cells for titer analysis. After 30 min incubation at 33°C, the virus was removed by washing and 5 mL of 1% agar/DMEM or 1% agar/DMEM containing 1 mM guanidine was added to the cells and incubated at 4 separate conditions: 33°C, 33°C and 1 mM guanidine, 39.5°C, and 39.5°C and 1 mM guanidine. After incubation for 2 days at 39.5°C or 4 days at 33°C, agar overlays were removed and cells stained with 0.1% crystal violet for plaque enumeration and calculation of PFU/mL. For RNA extraction and sequencing of progeny viruses/recombinants, prior to removal of agar overlay from plates incubated at 39.5°C and 1 mM guanidine, single plaques were picked and virus amplified on HeLa cells for 4 h. Infected cells were harvested and added to 1 mL TRIZOL for RNA extractions and RT-PCR. Sequences of RT-PCR products from nucleotides 4352 to 4955 in the poliovirus genome were generated using the UT Southwestern Sequencing Core.

## Author Contributions

Conceptualization, A.K.E, J.K.P.; Investigation, A.K.E, P.R.J.; Writing – Original Draft, A.K.E, J.K.P.; Writing – Review & Editing, A.K.E, P.R.J., J.K.P., S.E.W, M.J.M., A.N.; Funding Acquisition, J.K.P., S.E.W.; Resources, M.J.M., A.N., S.E.W.; Supervision, J.K.P.

## Acknowledgments

We thank Karla Kirkegaard for the Drug^R^/Temp^S^ (3NC-202gua^R^) and PV-2A144-DsRed poliovirus infectious clones, John Schoggins for the PV-2A144- GFP poliovirus infectious clone, and Lora Hooper and Lisa Forrest for providing bacterial strains. We also thank the Electron Microscopy Core Facility and the Flow Cytometry Core Facility at the University of Texas Southwestern Medical Center. Finally, we thank Matthew Taylor for a helpful tip regarding use of the Drug^R^/Temp^S^ poliovirus mutant.

This work was funded by NIH NIAID grants R01 AI74668 and R21 AI114927, and a Burroughs Wellcome Fund Investigators in the Pathogenesis of Infectious Diseases Award to J.K.P. The research of J.K.P. was supported in part by a Faculty Scholar grant from the Howard Hughes Medical Institute. Work in S.E.W.’s lab is supported by NIH NIAID grants R01 AI118807 and R21 AI128151.

## Supplementary Figure Legends

**Table S1.**
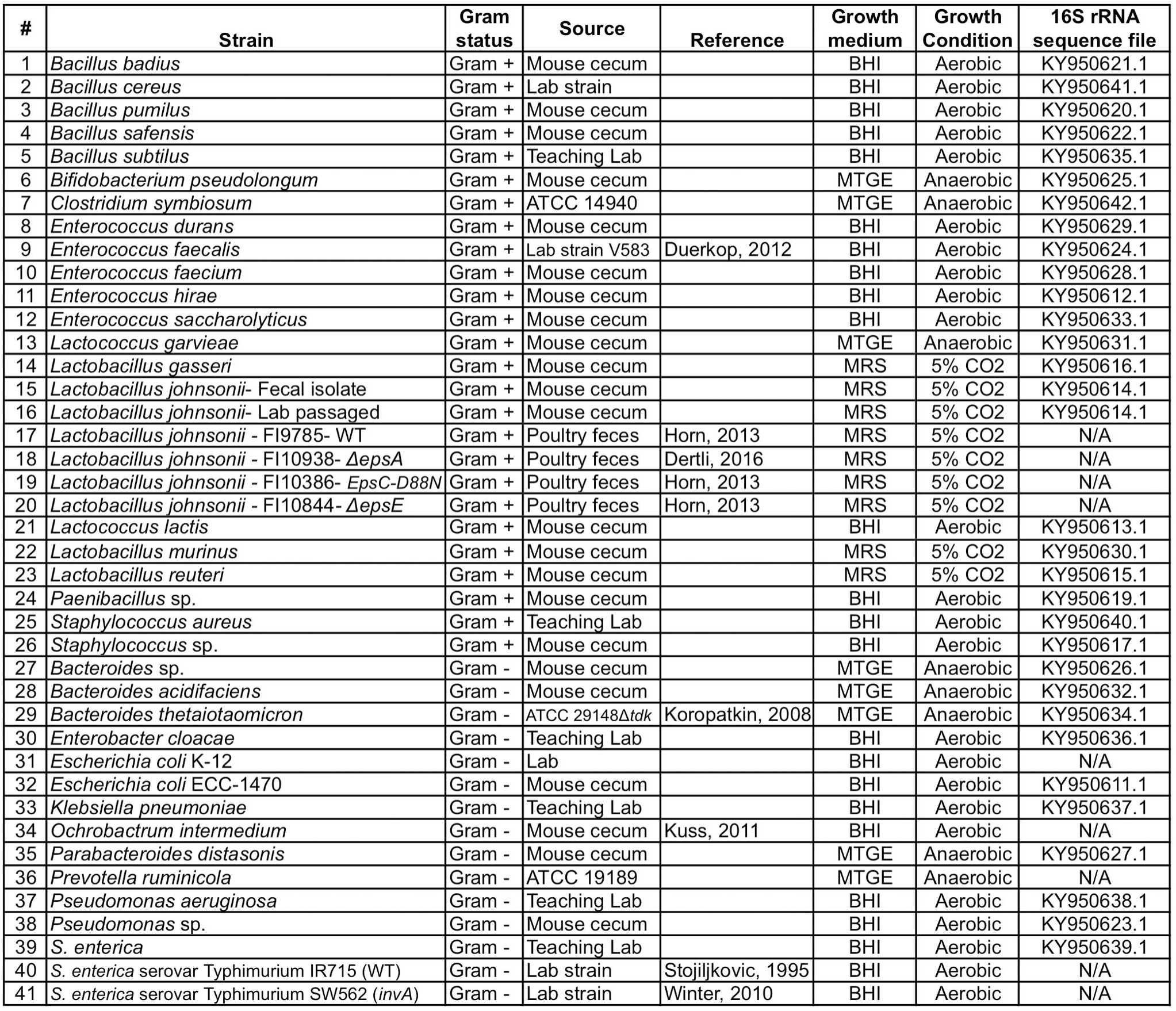
Related to Figure 1. Bacterial strains used in this study. Key: BHI= Brain-heart infusion medium, MTGE= anaerobic bacteria enrichment medium from Anaerobe Systems, MRS= deMan, Rogosa, and Sharp medium for culture of *Lactobacilli*. Link to all 16S rRNA sequences: https://www.ncbi.nlm.nih.gov/popset/?term=ky950611

**Figure S1.**
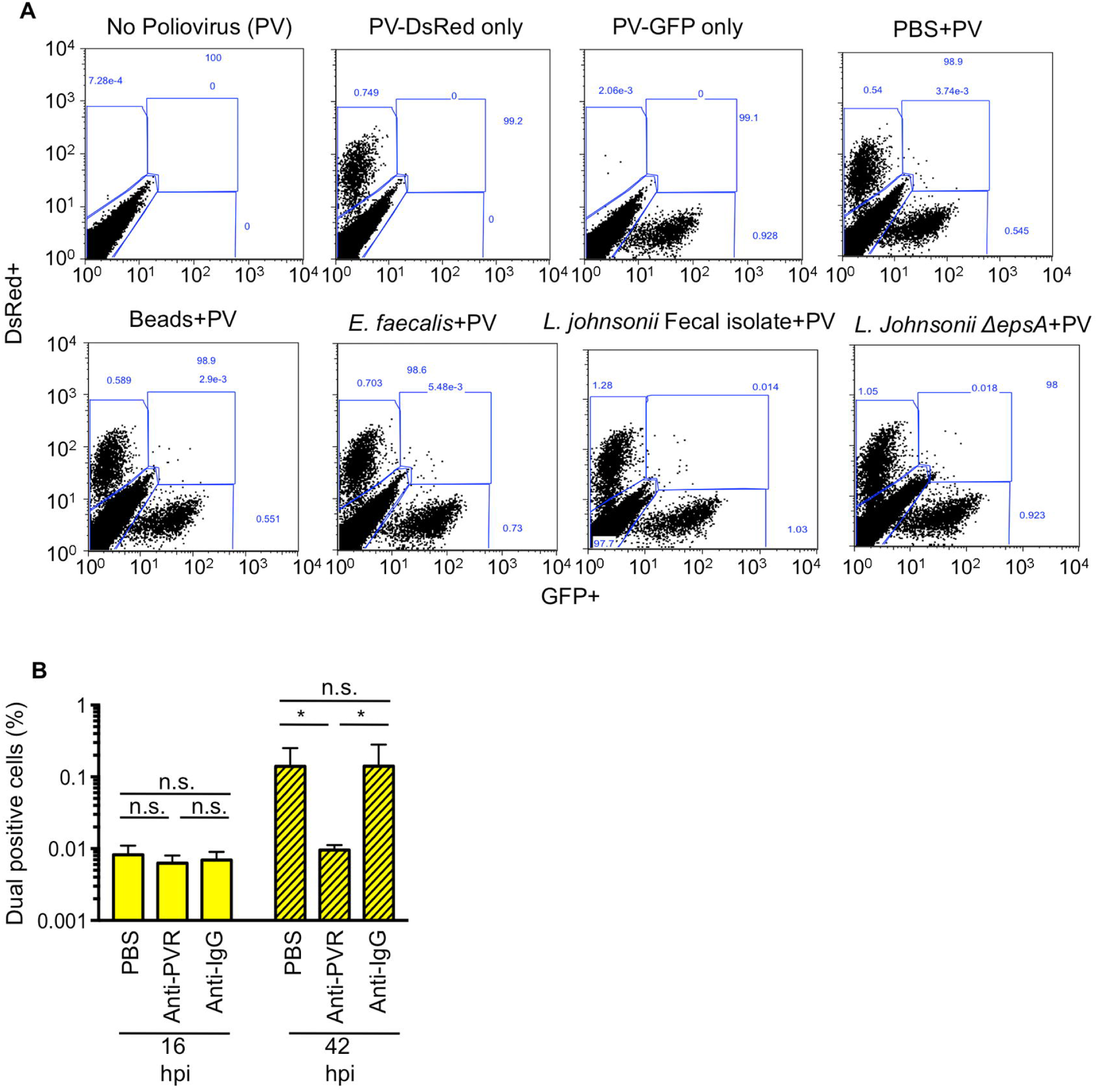
Related to Figures **3** and 4. Establishing a flow cytometry-based assay to quantify viral infection and co-infection. (A) Representative FACs diagrams showing gating strategies. In each experiment cells were mock infected or infected with 1x10^4^ PFU PV-DsRed and/or PV-GFP and positive gates were drawn from these samples and applied to all samples within that experiment. (B) Dual infected cells at 16 hpi are from infection with the initial inoculum and not secondary cycles of replication. 1x10^4^ DsRed- and GFP-expressing polioviruses were mixed prior to infection of 1x10^6^ HeLa cells (MOI of 0.01). HeLa cells were washed and media containing 6 μg of anti-PVR or anti-IgG antibody was added to cells. At 16 or 42 hpi cells were harvested and GFP+DsRed+ (dual positive cells) were quantified by flow cytometry. While the presence of PVR antibody reduced the number of dual positive cells at 42 hpi, likely due to co-infection during secondary cycles of replication, the number of dual infected cells at 16 hpi was unaffected. Data are shown as percentage of total cells counted and are mean ± SEM (n=6). \**p*<0.05 (Student’s unpaired t test).

**Figure S2.**
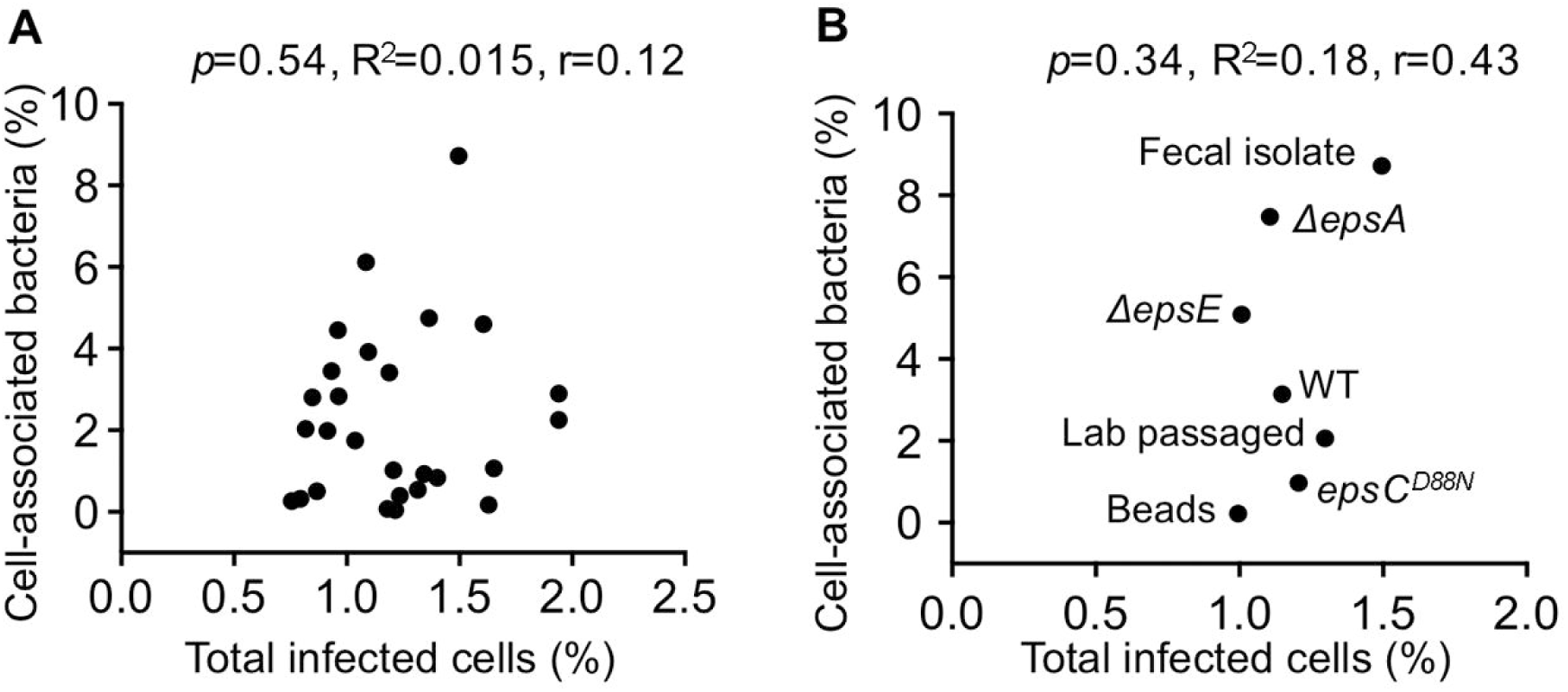
Related to Figure 5. The ability of bacteria to adhere to HeLa cells does not correlate with the total percentage of poliovirus-infected cells. (A) Scatter plot to test for correlation between total percent of infected cells and the percent of cell-associated bacteria for aerobic bacterial strains. Data points are the mean values presented in Figures 3C and 5B. *p*=0.54, R^2^=0.015, r=0.12 (Pearson’s correlation coefficient calculation). (B) Scatter plot to test for correlation between total percent of infected cells and the percent of cell-associated bacteria for *L. johnsonii* strains. Data points are the mean values presented in Figures 3E and 5D. *p*=0.34, R^2^=0.18, r=0.43 (Pearson’s correlation coefficient calculation).

**Figure S3.**
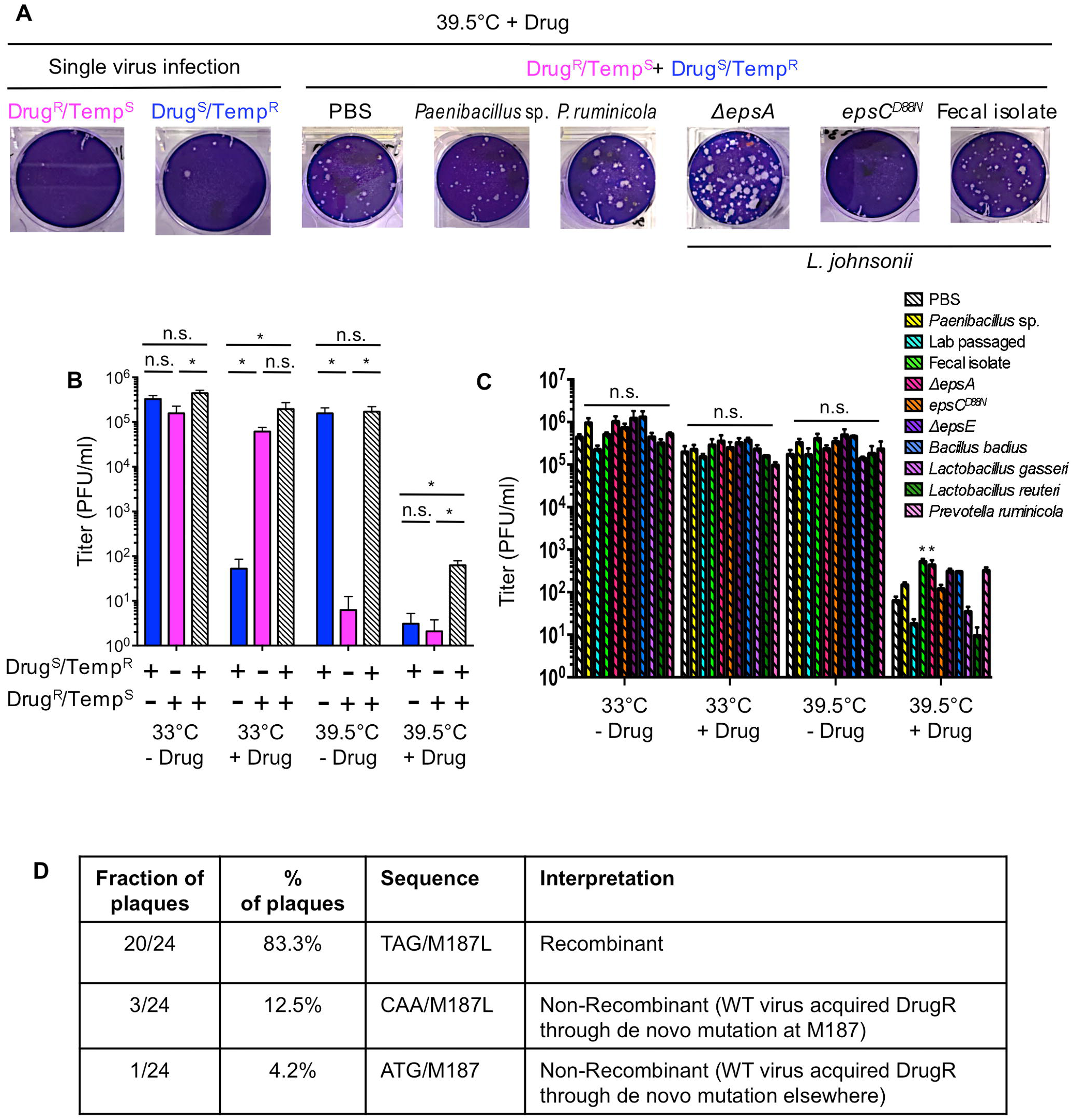
Related to Figure 6. Exposure to certain bacterial strains impacts poliovirus recombination frequency. (A) Representative plaque assays of recombination cross progeny viruses under nonpermissive growth conditions. 1x10^5^ PFU of each parental virus was used individually for infections (left) or mixed and incubated with 1x10^8^ CFU bacteria prior to infection of 1x10^7^ HeLa cells (right)(MOI of 0.01). Infection was allowed to proceed for 8 h under permissive conditions for both viruses (33°C without drug) and progeny viruses were quantified by plaque assay at both permissive and restrictive conditions to determine yields of individual parental viruses and recombinants. Plates from the 39.5°C+Drug group are shown. (B) Poliovirus yields (PFU/mL) at all growth conditions for progeny of infections with parental viruses alone or mixed together in PBS. Data are represented as mean ± SEM (n=6). \**p*<0.05 (one-way ANOVA followed by Dunnet’s multiple comparison test). (C) Poliovirus yields (PFU/mL) at all growth conditions of progeny viruses from dual parent infections (recombinant crosses) after exposure to bacteria. Data are represented as mean ± SEM (n=6). \**p*<0.05 versus PBS (one-way ANOVA followed by Dunnet’s multiple comparison test). (D) Sequence analysis of individual plaques picked from progeny of recombinant crosses. Plaques were picked from plates incubated at 39.5°C+Drug (see Fig. S3A), poliovirus was purified from the individual plaques, and the 2C region was sequenced from nucleotides 4352 through 4955 to determine whether the phenotypic recombinants were confirmed genotypically. The Drug^R^/Temp^S^ virus has a M187L amino acid change (via change from ATG to TAG at nucleotides 4682-4684) in the 2C protein that confers guanidine resistance as well as an additional V250A amino acid change (via change from GTC to GCG at nucleotides 4871-4873) in the 2C protein that contributes to the Temp^S^ phenotype (Kirkegaard and Baltimore, 1986). Recombination in the 190 nt interval between these two alleles of the parental viruses can generate M187L (Drug^R^)/V250 (Temp^R^) viruses. Therefore, we examined consensus sequences of this region in 24 plaques picked from plates incubated at 39.5°C+Drug to determine whether the predicted sequences were present. Of 24 plaques picked, all contained the wild-type allele at V250, indicating they were likely derived from the Drug^S^/Temp^R^ (wild-type) parental virus. Additionally, the majority (83.3%) had a TAG codon at nucleotides 4682-4684, confirming presence of the Drug^R^ allele from the Drug^R^/Temp^S^ parental virus. Therefore, 83.3% of phenotypic recombinants (Drug^R^/Temp^R^) were confirmed as genotypic recombinants. The remaining four plaques were likely not recombinants, but rather mutants generated by *de novo* acquisition of guanidine resistance, an expected phenomenon based on the error frequency of poliovirus: 3 plaques had CAA codons at M187, which generated the M187L amino acid change through *de novo* mutation, and 1 plaque had the wild-type ATG codon at M187, suggesting that a novel guanidine resistance mutation may have occurred elsewhere in the genome (Pincus et al., 1986).

